# Whole-brain modular dynamics at rest predict sensorimotor learning performance

**DOI:** 10.1101/2023.12.18.572267

**Authors:** Dominic I. Standage, Daniel J. Gale, Joseph Y. Nashed, J. Randall Flanagan, Jason P. Gallivan

## Abstract

Predictive biomarkers of cognitive performance are informative about the neural mechanisms underlying cognitive phenomena, and have tremendous potential for the diagnosis and treatment of neuropathologies with cognitive symptoms. Among such biomarkers, the modularity (subnetwork composition) of whole-brain functional networks is especially promising, due to its longstanding theoretical foundations and recent success in predicting clinical outcomes. We used functional magnetic resonance imaging to identify whole-brain modules at rest, calculating metrics of their spatio-temporal dynamics before and after a sensorimotor learning task on which fast learning is widely believed to be supported by a cognitive strategy. We found that participants’ learning performance was predicted by the strength of dynamic modularity scores (clarity of subnetwork composition), the degree of coordination of modular reconfiguration, and the strength of recruitment and integration of networks derived during the task itself. Our findings identify these whole-brain metrics as promising biomarkers of cognition, with relevance to basic and clinical neuroscience.

## Introduction

Resting-state functional magnetic resonance imaging (rsfMRI) is invaluable for investigating the brain’s instrinsic functional organisation, free from explicit task engagement (1, 2). Resting state functional connectivity (RSFC), determined by the pairwise covariation of haemodynamic signals at rest, has identified robust commonalities of this organisation across large sets of participants (3), while simultaneously providing measures for exploring individual and group differences (4, 5). These differences are the focus of intense research interest, due to their predictive relationships with a wide variety of cognitive measures (6–9), neuropathologies (10–15) and clinical interventions (16–18). As such, there is growing interest in RSFC measures that can account for behavioural variability across participant populations, with a view to identifying predictive biomarkers (4, 8, 12, 13, 16, 18, 19).

A promising trend in this body of work is the predictive value of resting-state whole-brain modularity, or the subnetwork composition of functional brain networks at rest (14, 20–22). Modularity conveys a number of advantages to complex systems, the gist of which is a balance between the competing demands of functional specialisation and integration. The brain is highly modular, both structurally and functionally, conveying benefits to distributed information processing and emergent dynamics (23) (see the Discussion). As such, its candidacy as a biomarker of cognitive performance is well founded.

Sensorimotor adaptation (SA) tasks are ideal for investigating individual differences in cognition, since participants show a high degree of variability in their use of cognitive strategies toward these tasks (24–26). On SA tasks, an established sensorimotor mapping is systematically altered, such as visual feedback of hand motion while reaching, so that participants are required to learn a new mapping for competent task performance. A large body of data supports the hypothesis that learning on SA tasks is driven by two components, one implicit and one explicit (27). The implicit component is slow, non-volitional, and widely believed to involve the recalibration of an internal model encoded by the motor system (28, 29). The explicit component is fast, volitional and corresponds to the use of a strategy toward the task, such as aiming to the left of a bullseye to counteract a strong rightward wind (30–32).

We recently studied the whole-brain dynamics of SA, finding that cohesive or coordinated modular reconfiguration was associated with fast learning during the task, and by association, with the use of a cognitive strategy. We also derived task-based functional networks from the dynamic modules, finding subnetworks that were associated with fast learning during early adaptation and re-adaptation respectively (26). Here, we refer to these networks as ‘the learning network’ and ‘the relearning network’ because of these behavioural associations. Importantly, our in-scanner SA task was preceded (and followed) by a resting scan, enabling us to investigate the extent to which spontaneous patterns of brain activity at rest are predictive of subsequent task performance.

We tested three principal hypotheses on resting-state biomarkers of fast learning, and by association, cognitive performance. We hypothesised that fast learning among healthy young adults would be predicted by the degree of (1) dynamic modularity, (2) coordinated modular reconfiguration, and (3) recruitment and integration of task-derived networks. Hypothesis 1 was based on the aforementioned findings that static modularity at rest can predict training-based improvements in cognitive performance among patient groups (20) and healthy older adults (21, 22). Hypothesis 2 was based on our earlier finding that coordinated modular reconfiguration is ‘conducive’ to a cognitive strategy toward adaptation, but does not directly implement that strategy (26). We reasoned that this conduciveness might also be observable at rest, prior to the task. Hypothesis 3 was based on our earlier finding that the learning and relearning networks are associated with fast adaptation. We reasoned that if recruitment of these (or other) task-based networks are supportive of task performance [and/or individual differences in performance (26)] then a predisposition toward their composition would be advantageous. Conversely, we reasoned that if their integration is disruptive to task performance (26) then such a predisposition would be disadvantageous. All predictive measures were re-quantified after the task in a second resting scan, enabling us to determine whether participants’ intrinsic whole-brain networks were modified by adaptation, as seen with some other tasks and network measures (33–35).

## Results

We analysed rs-fMRI data from a study (26) in which participants performed a classic visuomotor rotation (VMR) task (36), controlling a small force sensor with the hand in order to move a cursor to a target that appeared randomly at one of eight locations around an invisible circle. On each of two testing days (separated by 24 hours) participants performed a baseline block, during which motion of the cursor was veridically mapped to motion of the hand, and a learning block, during which the correspondence between hand movement and cursor movement was rotated clockwise by 45 degrees. This rotation required participants to adapt their movements (in the counterclockwise direction) to intercept the target. Resting scans were taken immediately before and after the task, during which participants lay still with their eyes open, maintaining central fixation. See Methods for details of the full experimental regimen.

Participants’ group-level behaviour was typical of this class of task, as (on average) they reduced their errors during the learning block and also showed ‘savings’ or faster relearning upon re-exposure to the VMR on the second day (Figure 2A). However, these group-level data obscured significant inter-participant variability (Figure 2B), as some participants were fast learners on each day (fast-fast, FF), others were slow learners on each day (slow-slow, SS) and others still were fast learners on the first day and slow learners on the second day (slow-fast, SF). To determine if these individual differences revealed a continuum of learning outcomes or a limited number of learning profiles, we clustered (see Methods)) participants’ early error on each day, late error on each day, and savings (see caption to Figure 2). These measures are commonly used to characterise behaviour on VMR tasks (25, 26, 31, 32), as they capture participants’ rate of learning, completeness of learning, and improvement in learning rate over the two days, respectively.

Our clustering revealed three distinct subgroups of learners, well described as FF, SS and SF (Figure 2B). Notably, the FF and SS subgroups clearly correspond to more explicit and implicit learners respectively, whereas the SF subgroup is ambiguous in this regard (26). Due to issues of statistical power relating to the subgroup sizes [N(FF) = 15, N(SS) = 10, N(SF) = 7] we sought to identify a single scalar measure corresponding to subgroup membership, enabling us to investigate brain-behaviour relationships with standard correlation methods (prior to post-hoc testing for subgroup differences). We therefore ran principal component analysis on the five behavioural measures used to identify the subgroups. The first component (PC1) accounted for 61% of the variance in the data and accurately classified 31 out of 32 participants (Figure 2C). The interested reader can find the full behavioural analysis in Standage et al. (26).

To test our hypotheses on dynamic modularity at rest and its efficacy as a predictor of upcoming task performance, we constructed ‘multi-slice’ functional networks from the covariation of regional brain signals (in sliding time windows) during a resting scan immediately before (and after) the task. We partitioned these resting-state networks into spatio-temporal modules that maximized a quality function *Q* (37) and verified that participants’ whole-brain networks showed significant modularity (38). Thus, we established that participants’ intrinsic functional networks were composed of interacting subnetworks before and after task performance (Supplementary Materials Figure 1).

**Fig. 1.**
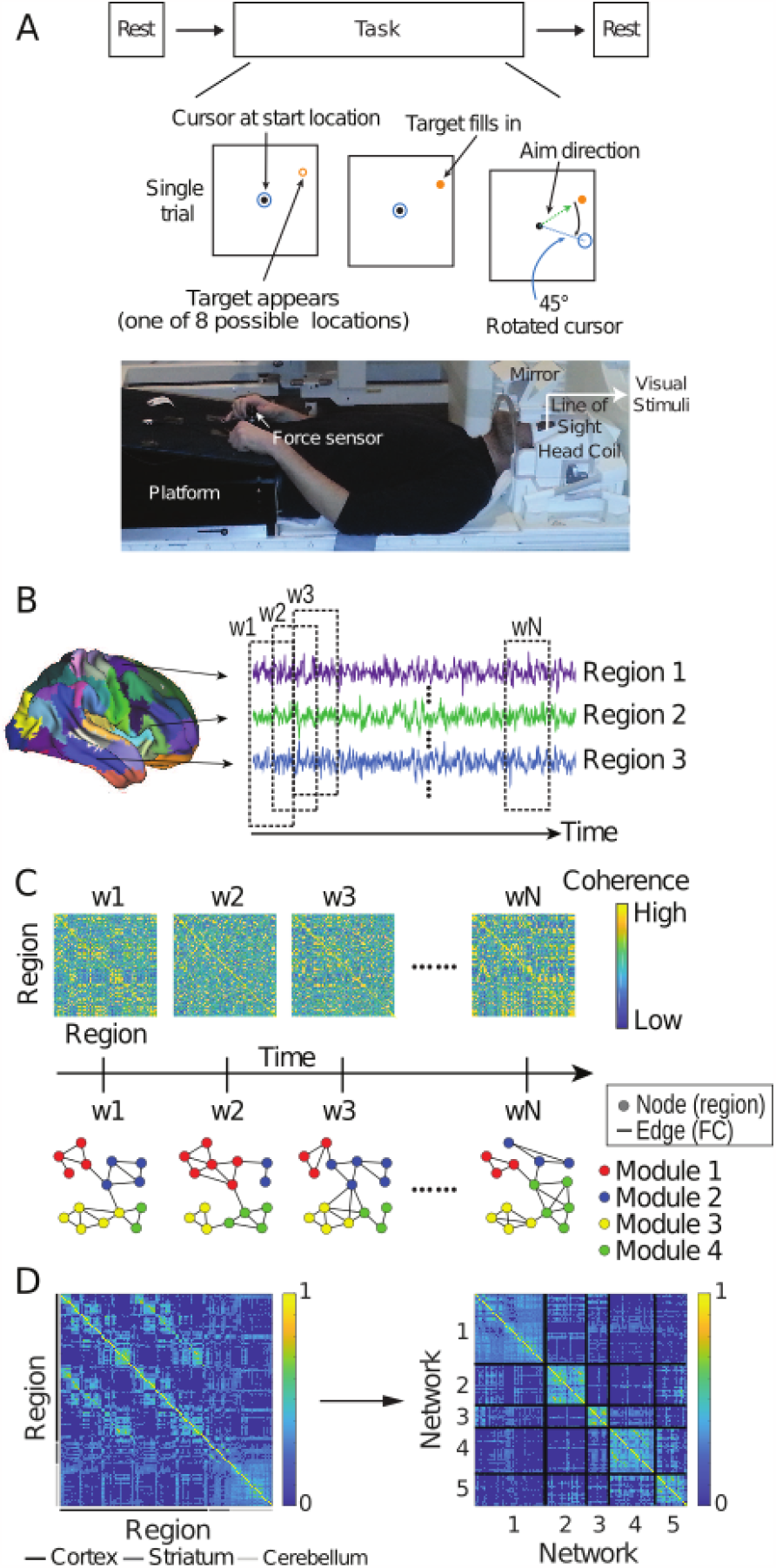
Overview of the task and neural analyses. **(A)** Participants underwent a resting scan before and after a visuomotor rotation (VMR) task, during which the viewed cursor, controlled by the hand, was rotated about the movement start location. **(B)** For each participant, the cerebral cortex, striatum, and cerebellum were parcellated into discrete regions and the average %BOLD time series was extracted from each region during resting and task scans (3 example cortical regions are shown). **(C)** The coherence of (Haar family) wavelet coefficients was calculated for each pair of regions in sliding time windows to construct functional connectivity matrices for each window (w1 – wN, see Methods). Time-resolved clustering methods were applied to the resulting multi-slice networks, identifying dynamic modules across time slices (4 modules in the schematic). **(D)** Module allegiance matrix (left) showing the probability that each pair of brain regions was in the same module during early learning, calculated over all participants, modular partitions and time windows (left). Arrow depicts the clustering of this matrix, identifying networks (right) that summarise the modular dynamics (see Methods).

**Fig. 2.**
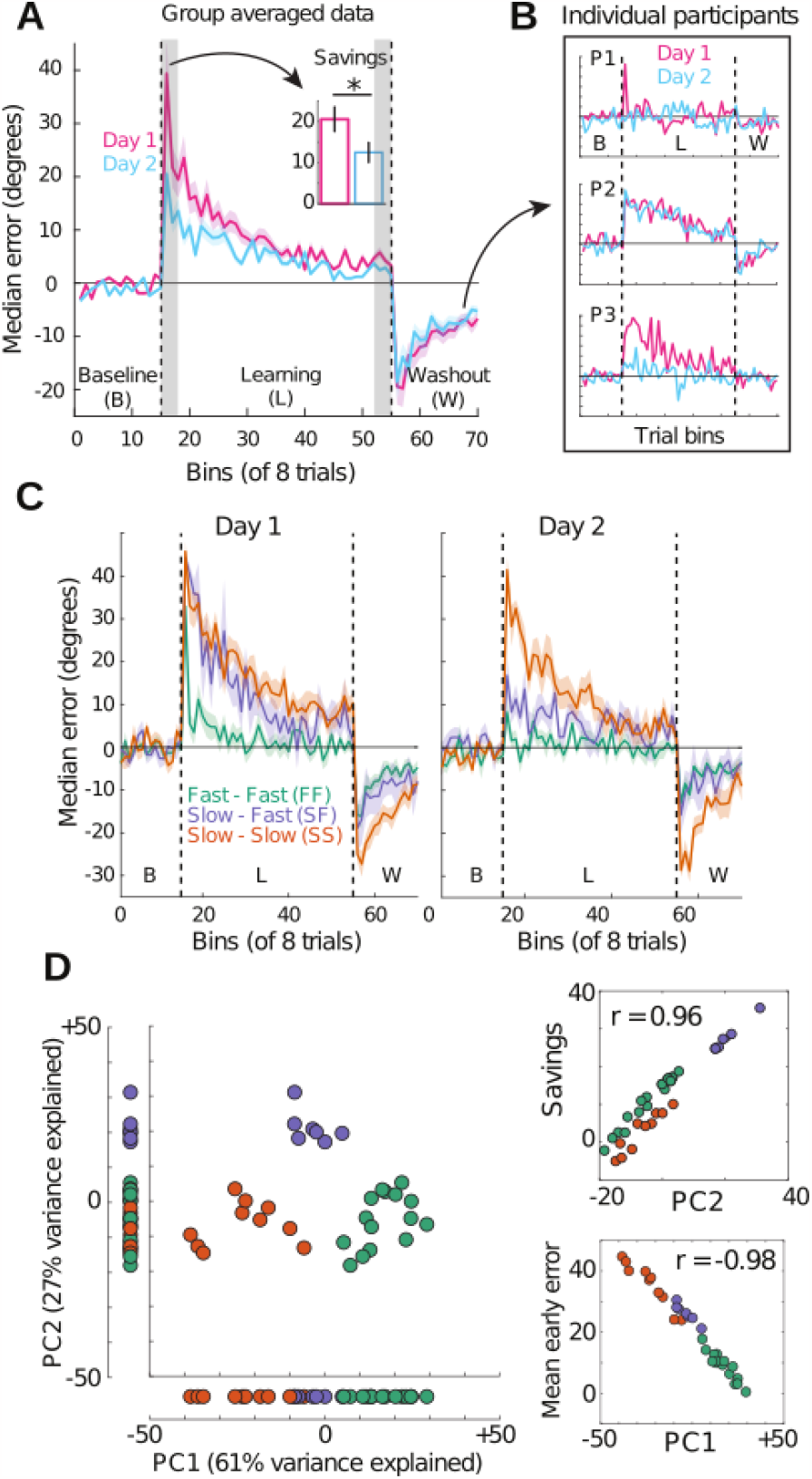
Participants were assigned to one of three subgroups by a clustering of their behavioural data. **(A)** Mean (across participants) of bin median errors during baseline (non-rotation), learning (45 deg rotation of the cursor) and ‘washout’ (non-rotation, to unlearn the mapping prior to the second day) on Day 1 (pink) and Day 2 (blue). Bins consisted of 8 consecutive trials, during which the target was chosen at random without replacement from 8 equidistant locations around an invisible circle. Early and late error were defined as the first and last three learning bins respectively (grey shading). Ribbons show standard error (*±*1 SE) and the dashed vertical lines demark the 3 task blocks. Savings (inset, early error on Day 1 minus early error on Day 2) was significant at the group level (paired t-test; t(31) = 6.122, P = 8.666e-7). **(B)** The group-averaged approach in ‘A’ obscures individual differences in learning. The trajectory of errors by three example participants is well described as fast-fast (FF, top, fast on Day 1 and fast on Day 2), slow-slow (middle, SS, slow on Day 1 and slow on Day 2) and slow-fast (SF, bottom, slow on Day 1 and fast on Day 2). **(C)** A clustering of participants’ early error on each day, late error on each day and savings (5 variables in total) identified three subgroups of participants whose mean bin median errors resemble the example participants in ‘B’ on Day 1 (left) and Day 2 (right). **(D)** Top two PCA components for the 5 learning measures across participants. Data points correspond to participants, color-coded by their cluster-assigned subgroup (legend in panel C). The horizontal axis shows that PC1 accurately classifies 31 of 32 participants (97% accuracy). Scatter plots on the right show that PC1 and PC2 closely correspond to mean early error across days and savings, respectively.

**Fig. 3.**
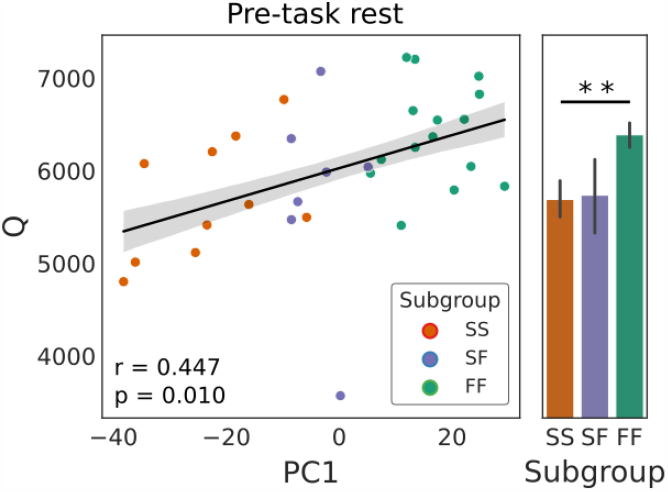
Prior to learning, greater dynamic modularity at rest predicted better task performance. The scatter plot shows the mean quality function score Q for each subject (quantifying modularity, see text) against PC1, our proxy for behavioural subgroup membership. Filled circles correspond to the FF, SS, and SF behavioural subgroups (see legend). The fitted line shows the best linear fit, where the shaded area shows *±*1 standard error (SE). Subgroup means *±*1 SE are shown as bar graphs to the right of the scatter plots. Q was significantly greater among FF participants than SS [t(23) = 2.931, p = 0.008] but did not differ statistically between FF and SF [t(20) = 1.907, p = 0.071] or between SF and SS [t(15) = 0.109, p = 0.914]. The two stars in the bar plot indicate statistical significance (*P <* 0.01).

### Dynamic modularity at rest predicted learning performance

To test our first hypothesis that greater modularity at rest would predict a faster learning profile over the two-day task, we averaged the quality function score *Q* (37) over all modular partitions for each participant (see Methods). We then calculated the correlation between PC1 and this mean *Q* score (henceforth *Q*). Consistent with our hypothesis, the correlation was indeed significant (Pearson’s r = 0.447, p = 0.010), where *Q* was significantly greater among FF participants than SS, but did not differ statistically between either of these subgroups and SF. Thus, greater dynamic modularity of functional networks at rest was associated with faster learning, and by association, a greater contribution of cognitive strategies to learning.

### Coordinated modular reconfiguration at rest predicted learning performance

To test our second hypothesis that the coordination of modular reconfiguration at rest would predict participants’ learning profiles, we calculated the cohesive and disjointed flexibility (a.k.a. cohesion strength and disjointedness) of participants’ brain regions during the first resting scan (see Methods). Cohesion strength refers to the number of times a given region changes modules together with each other region, summed over all other regions. In contrast, disjointedness refers to the number of times a given region changes modules on its own, relative to the number transitions between time windows (the number of time windows minus one) (39). For each participant, we therefore calculated each of these measures for each brain region in each partition, before averaging all partitions and regional scores (see Methods).

Consistent with our hypothesis, cohesion strength was positively correlated with PC1 (r = 0.425, p = 0.015) and was significantly stronger by the FF subgroup than the SS subgroup, but did not differ statistically between either of these subgroups and SF (Figure 4A). Inversely, disjointedness was negatively correlated with PC1 (r = -0.431, p = 0.014) and was significantly stronger by the SS subgroup than FF, but did not differ statistically between either of these subgroups and SF (Figure 4B). As was the case during early learning (26), both measures captured global, whole-brain dynamics, as the correlation between cohesion strength and PC1 was positive for 138 out of 142 brain regions, whereas the correlation between disjointedness and PC1 was negative for 112 out of 142 regions (Figure 4C). Thus, coordinated modular reconfiguration is a whole-brain property that is not only conducive to cognitive performance during SA (26), but is also predictive of future performance.

**Fig. 4.**
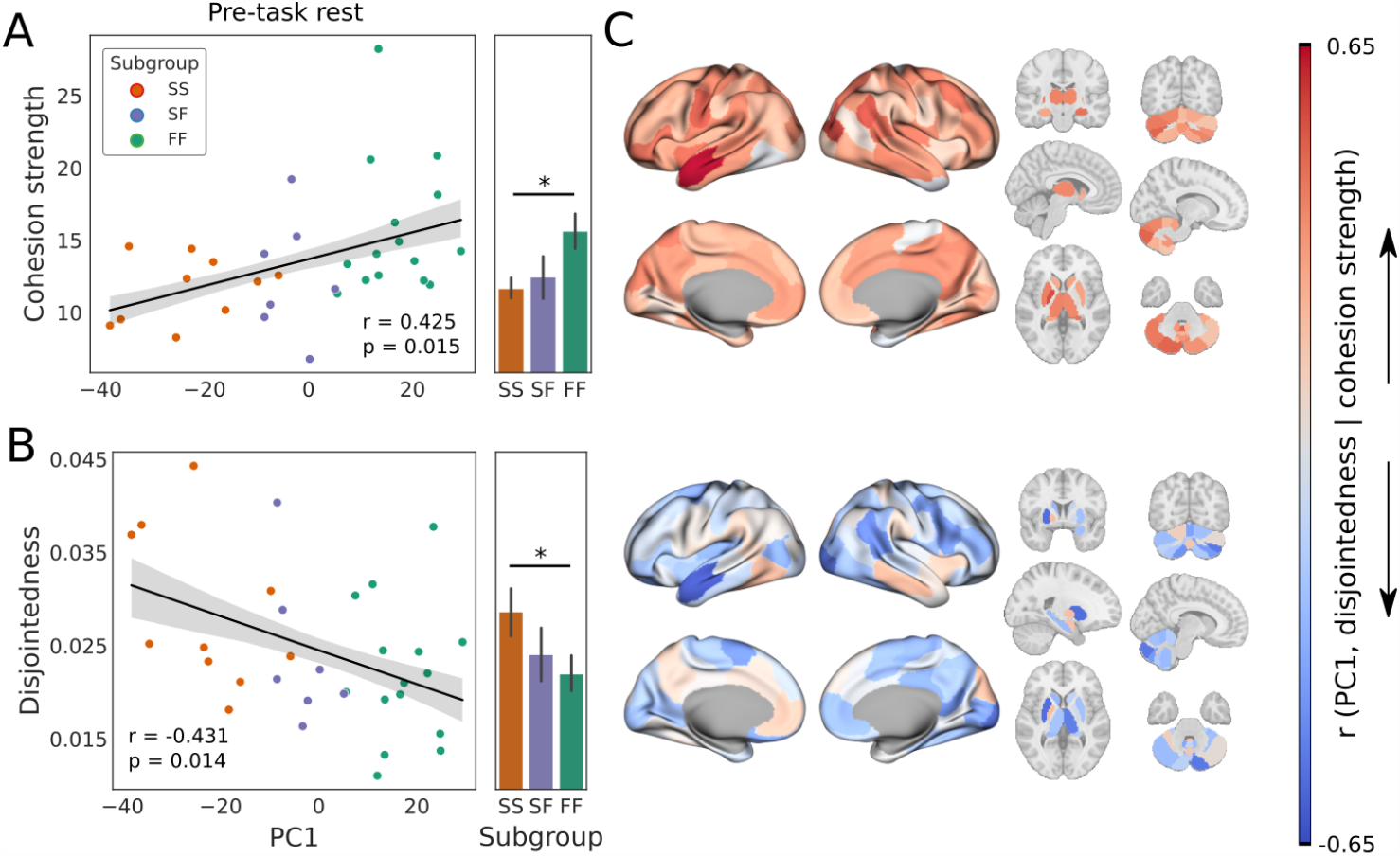
Coordinated (A) and uncoordinated (B) modular reconfiguration at rest predicted fast and slow learning profiles on the upcoming task, respectively. **(A)** Mean cohesive flexibility (cohesion strength, see text) as a function of PC1. FF was more cohesive than SS [t(23) = 2.518, p = 0.019], but SF did not differ significantly from FF [t(20) = 1.556, p = 0.135] or SS [t(15) = 0.515, p = 0.614]. **(B)** Mean disjointed flexibility (disjointedness, see text) as a function of PC1. SS was more disjointed than FF [t(23) = -2.089, p = 0.048], but SF did not differ significantly from SS [t(15) = -1.115, p = 0.282] or FF [t(20) = -0.596, p = 0.558]. **(C)** Brain plots show correlations between PC1 and cohesion strength (upper plots) and disjointedness (lower plots) for each region. Correlation values associated with cohesion strength (mostly positive) and disjointedness (mostly negative) are shown with a divergent color scheme, ranging from strongly negative (dark blue) to strongly positive (dark red). In bar plots, stars indicate significant differences (*p <* 0.05).

### Resting-state recruitment and integration of task-based networks predicted learning performance

To test our third hypothesis that the recruitment of task-relevant networks at rest would predict participants’ learning profiles, we assigned each brain region to one of five task-based networks derived during early learning (26), when the use of cognitive strategies toward the task has been shown to be most pronounced [supporting the fastest learning (30, 40)]. The ‘recruitment’ of an individual network can be measured in terms of the ‘interaction’ between two networks by calculating the network’s interaction with itself (41). Under this approach, the interaction between two networks is defined by *I*_*k*1,*k*2_ = (∑*i∈C*_*k*1_, *j∈ C*_*k*2_*P*_*i,j*_)*/* (|*C*_*k*1_| | *C*_*k*2_|), where *C*_*k∈{*1,2}_ are modules |*C*_*k*|_ is the number of regions they contain, and *P*_*i,j*_ is a ‘module allegiance matrix’ quantifying the probability that each pair of regions was in the same module (here, during the first resting scan). Under this approach, recruitment is therefore measured by letting *k*1 = *k*2, quantifying the consistency of regional interactions within a specified network over time.

Following false discovery rate (FDR) correction over the five networks (see Methods), we found a significant positive correlation between recruitment of the relearning network and PC1 (r = 0.501, p = 0.003, adjusted p = 0.017). This finding was surprising to us, since the relearning network was not associated with participants’ behavioural profiles during the task until the *second* day of the experiment (26). As was the case with *Q* and modular reconfiguration above, the FF and SS subgroups were significantly different according to this measure (FF recruited the network more strongly than SS) but neither of these subgroups differed statistically from SF (Figure 5A).

**Fig. 5.**
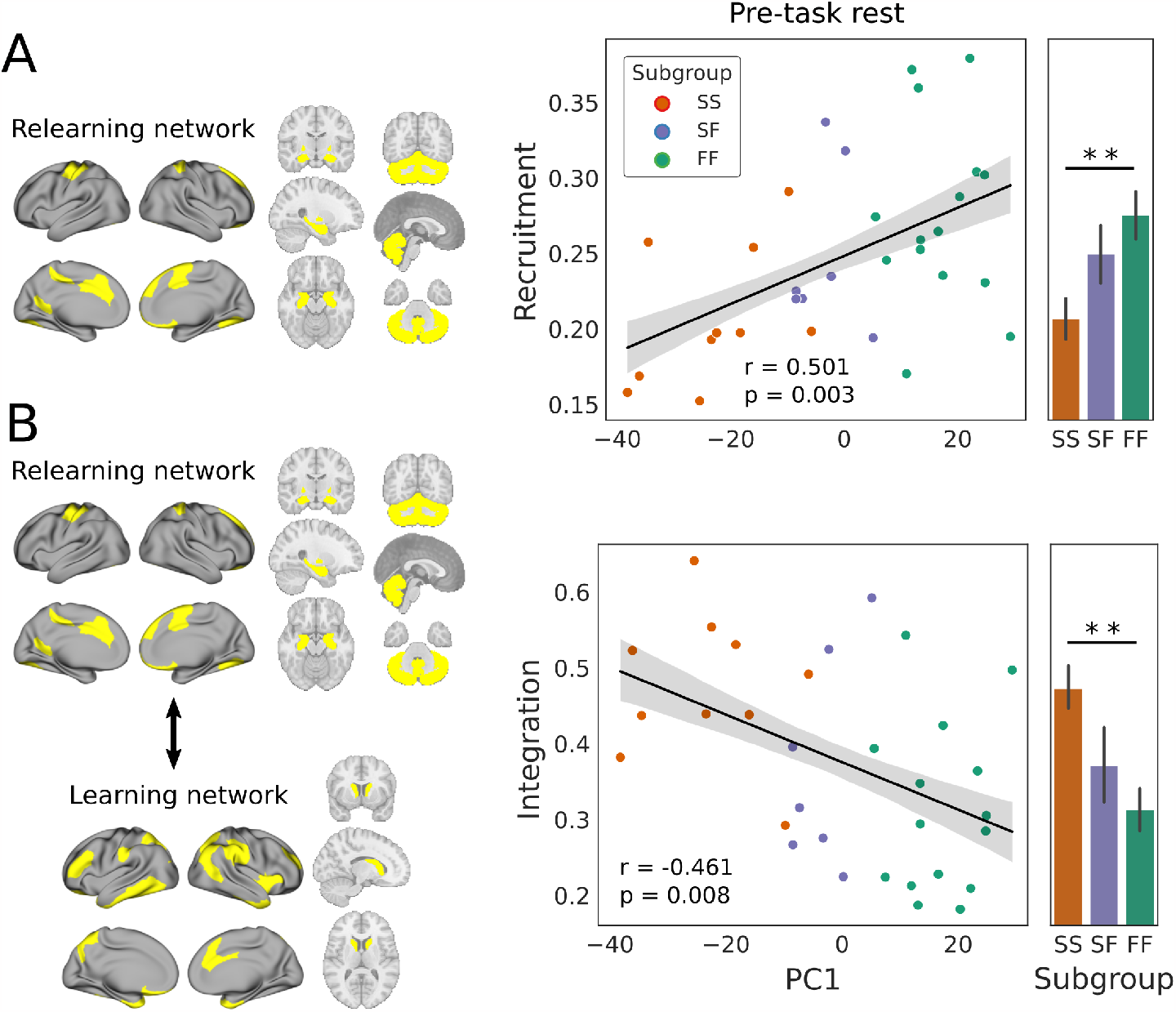
(A) Prior to learning, stronger recruitment of the relearning network, and weaker integration between the learning and relearning networks, predicted faster learning and thereby a more strategic (cognitive) approach to the task. The relearning network (left) consisted of regions spanning contralateral motor cortex, bilateral cerebellum, medial prefrontal cortex and several subcortical structures (bilateral hippocampus, pallidum, amygdala and accumbens). Recruitment of this network at rest was positively correlated with PC1, where recruitment by FF was stronger than SS [t(23) = 3.028, p = 0.006], but recruitment by SF did not differ statistically from that of FF [t(20) = 1.013, p = 0.323] or SS [t(15) = 1.858, p = 0.083]. **(B)** The learning network (left) consisted of regions spanning the anterior temporal pole, inferior and superior parietal, dorsolateral prefrontal cortex and the bilateral caudate. Its pre-task integration with the relearning network was negatively correlated with PC1, where integration was stronger by the SS subgroup than FF [t(23) = 3.574, p = 0.002] but did not differ statistically between SF and FF [t(20) = 1.061, p = 0.301] or SS [t(15) = -1.673, p = 0.115]. Error bars show ±1 SE. In bar plots, two stars indicate significant differences (*p <* 0.01).

Next, we sought to determine whether integration of the relearning network with any of the other four task-based networks would be predictive of participants’ behavioural profiles. The integration between two networks *k*1 ≠ *k*2 can be measured by the normalised interaction between them 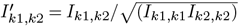 (41). As such, integration quantifies the degree to which networks interact over time, relative to their size. Following FDR correction over the other four networks (see Methods), we found a significant negative correlation between integration of the learning and relearning networks and PC1 (r = 0.461, p = 0.008, corrected p = 0.032). These two networks were significantly more integrated among the SS subgroup than FF, but neither of these subgroups differed statistically from SF (Figure 5B).

Because we measured recruitment and integration at rest, prior to learning, and the five networks underlying these measurements were derived during learning itself, we wondered if our results would hold if summary networks were derived at rest. In this regard, we reasoned that if pre-task recruitment conveyed a performance advantage on the actual task (as evidenced by its correlation with PC1) then the recruited network should be sufficiently distinguishable to be derived before the task. We therefore applied our network-derivation procedure (26) to the first resting scan. Notably, the procedure identified four networks, not five (see Supplementary Materials Figure 2), implying that modular reconfiguration during the task entailed the construction of new, task-appropriate functional networks. We then identified the network derived during rest that was most similar to the relearning network (derived during early learning) where similarity was determined by the Jaccard index (42), *i*.*e*. the similarity of two networks was quantified as the size of their intersection divided by the size of their union (multiplied by 100). We then calculated the recruitment score for the most similar resting-state summary network (Jaccard index = 76%). Crucially, the results of this analysis replicated the recruitment results above. That is, during the first resting scan, we found a significant positive correlation between the recruitment of our ‘resting-state relearning network’ and PC1 (r = 0.500, p = 0.004, corrected p = 0.014), where recruitment was stronger by the FF subgroup than SS [t(23) = 2.918, p = 0.008], but did not differ significantly between SF and FF [t(20) = 0.503, p = 0.621] or SS [t(15) = 1.724, p = 0.105].

These results not only indicate that the regions of the relearning network were sufficiently coupled during pre-task rest to be distinguishable as a network at that time, but they also confirm that recruitment of this resting-state network was predictive of subsequent learning performance. For completeness, we calculated the recruitment score for this network during early relearning on Day 2 of the task, when the relearning network distinguished the FF and SF subgroups from the SS subgroup in our earlier study (26). Once again, we replicated our results, as we found a significant positive correlation between recruitment and PC1 (r = 0.450, p = 0.010, corrected p = 0.039), where recruitment was stronger by the FF [t(23) = 2.412, p = 0.024] and SF [t(15) = 3.179, p = 0.006] sub-groups than the SS subgroup, but did not differ significantly between FF and SF [t(20) = -0.157, p = 0.877]. This stringent test shows that this network was not only predictive of performance on the upcoming task, but also, that it could ‘stand in’ for the relearning network during relearning itself.

Following these results, we sought to determine if there was a significant correlation between PC1 and the integration of the above ‘resting-state relearning network’ with a ‘resting-state learning network’. The latter, however, appears to be highly task-specific, as all four resting-state networks were at least 59% similar to one of the other four task-based networks, but no single resting state network was more than 20% similar to the learning network derived during adaptation. At first glance, this finding may appear quizzical, in light of our earlier result that integration of the learning and relearning networks during rest predicts slow learning (Figure 5B). There is nothing quizzical about it, however. We assigned task-based network labels (from 1 to 5) to brain regions and then calculated the recruitment and integration of each group of labelled regions at rest. It turns out that the learning network had not been ‘put together’ prior to the task. This finding indicates that slower learning was actually predicted by greater integration of its *dispersed* regions with the relearning network. In other words, the learning network was not recruited until the task was performed, but we were still able to calculate the integration of its regions (distributed across multiple networks) with the relearning network. This measure of integration predicted slow learning (Figure 5B).

### Resting-state recruitment and integration of task-based networks differed before and after learning

Having tested our hypotheses on predictive biomarkers of cognition, we sought to determine whether resting-state modular dynamics would differ before and after task performance. That is, we asked whether resting-state modular dynamics were altered by learning. To answer this question, we calculated whole-group differences (Rest 2 minus Rest 1) for the five modularity-based measures taken above, *i*.*e*. for Q, cohesion strength, disjointedness, recruitment of the relearning network, and the integration of the relearning network with (the dispersed regions of) the learning network. Following FDR-correction over the five tests, none of these differences were significant [paired t-test, Q: t(31) = 0.880, p = 0.386, corrected p = 0.482; cohesion strength: t(31) = -0.226, p = 0.823, corrected p = 0.823; disjointedness t(31) = 0.953, p = 0.348, corrected p = 0.482; recruitment: t(31) = 2.420, p = 0.022, corrected p = 0.054; integration: t(31) = 2.732, p = 0.010, corrected p = 0.052].

We also calculated the correlation between PC1 and each of these measures during the second resting scan, determining if the correlation was different than during the first resting scan (Steiger’s method for dependent correlations). Following FDR-correction over the five tests, the correlation between PC1 and recruitment of the relearning network was significantly different (Figure 6A). We therefore calculated the difference in recruitment for each subgroup, finding a significant increase during post-task rest among SS participants, where this increase was significantly greater than the change in recruitment by FF participants (Figure 6B). The correlation between PC1 and integration was also significantly different following FDR-correction (Figure 6C) where we found a significant decrease during post-task rest among SS participants, the magnitude of which was significantly greater than the change in integration by FF and SF participants (Figure 6D). Thus, these changes in intrinsic network composition were driven by the SS subgroup, who recruited the relearning network more strongly in the second resting scan, relative to the first, and disentangled it from the dispersed regions that formed the learning network during the task. However, this reorganisation does not appear to have influenced the behaviour of SS participants on the second day. In other words, resting-state recruitment and integration of the learning and relearning networks by SS became more like those of FF and SF after learning, but these changes did not predict improved performance the next day. We return to these findings in the Discussion.

**Fig. 6.**
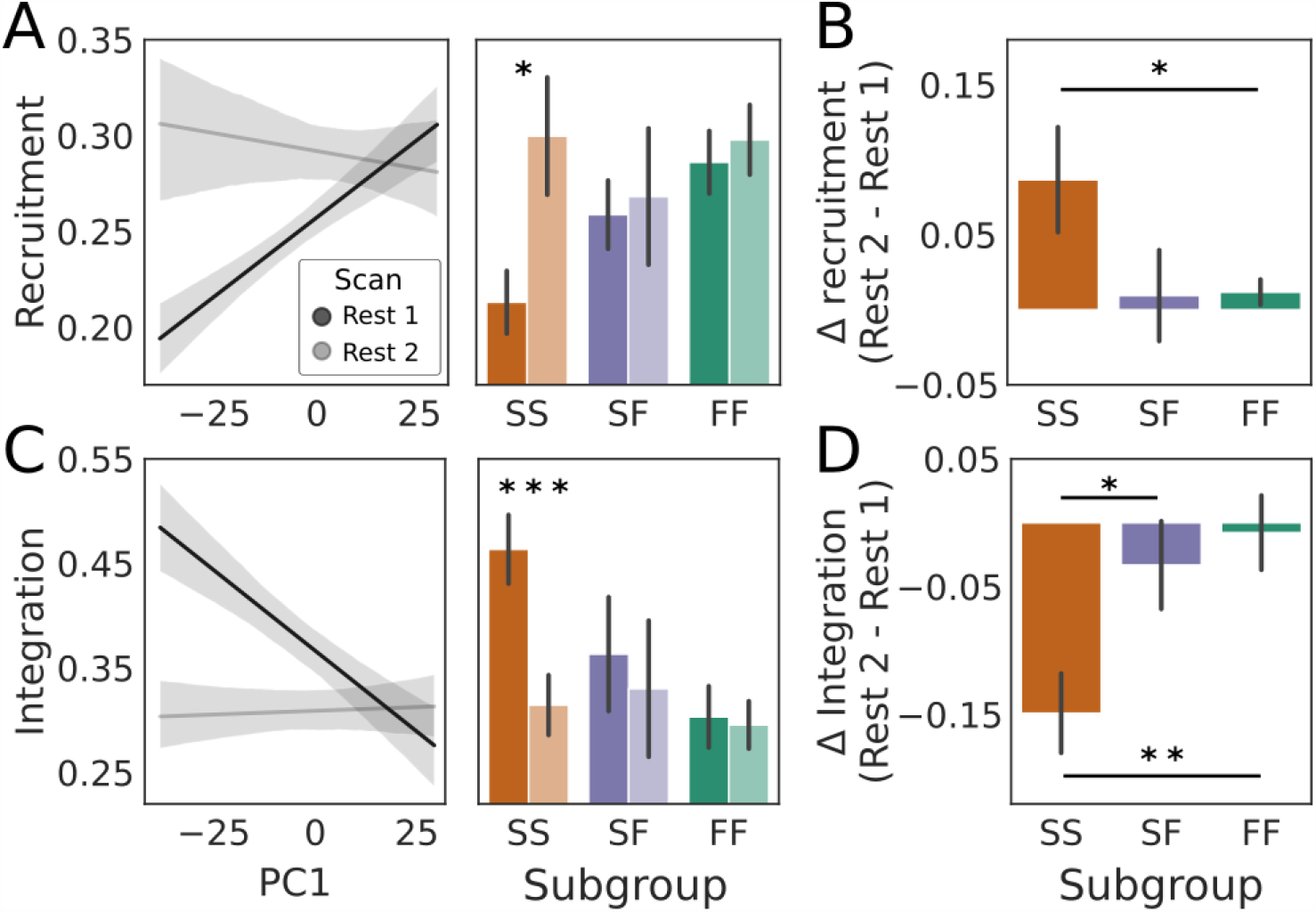
**A** The correlation between PC1 and recruitment of the relearning network was stronger [Steiger’s test for dependent correlations: t(30) = 3.636 p = 0.001, corrected p = 0.005] during the first (pre-task) resting scan (Rest 1, black: r = 0.501, p = 0.003, per Figure 5A) than the second (post-task) resting scan (Rest 2, grey: r = -0.087, p = 0.635). Shading corresponds to *±*1SE. The change in recruitment was driven by the SS subgroup (right panel), whose recruitment was significantly greater [paired t-test: t(9) = 2.475, p = 0.035] during the second resting scan (lighter shade) than the first (darker shade). Neither of the FF [t(14) = 1.386, p = 0.187] or SF [t(6) = 0.311, p = 0.766] subgroups differed significantly between resting scans. **(B)** The change in recruitment by SS was significantly greater than that by FF [two-sample t-test: t(23) = -2.487, p = 0.021] but did not differ significantly between SS and SF [t(15) = 0.096, p = 0.924] or between SF and FF [t(20) = -1.573, p = 0.137]. **(C)** The correlation between PC1 and integration (of the relearning network with the regions of the learning network) was weaker [Steiger’s test for dependent correlations: t(30) = -3.136, p = 0.004, corrected p = 0.010] during the first resting scan (Rest 1, black, r = -0.446, p = 0.010, per Figure 5B) than the second resting scan (Rest 2, grey, r = 0.025, p = 0.890). The change was again driven by the SS subgroup (right panel), whose integration was significantly weaker [t(9) = -4.798, p = 9.758e-4] during the second resting (lighter shade) scan than during the first (darker shade). The FF [t(14) = -0.266, p = 0.794] and SF [t(6) = -0.964, p = 0.372] subgroups did not differ significantly between resting scans. **(D)** The change in integration by SS was significantly greater than that by SF [two-sample t-test: t(15) = -2.468, p = 0.027] and FF [t(23) = 3.230, p = 0.004] but FF and SF were not significantly different [t(20) = 0.520, p = 0.609]. In all panels, stars indicate statistical significance (one star: p < 0.05; two stars: p < 0.01; three stars: p < 1e-3).

### The organisation of dynamic functional connectivity was more similar between resting scans than between either resting scan and learning

Our derivation of four summary networks from pre-task rest (compared to the five networks derived from early learning) suggests a fundamental difference between the organisation of dynamic, wholebrain functional networks at rest and during the task. In light of this finding, we predicted that the module allegiance matrices from which the networks were derived would be more similar when the resting scans were compared to each other than when either resting scan was compared to early learning. To test this prediction, we calculated the pairwise correlations between module allegiance matrices from these three scans. All three correlations were strong [Rest 1 (pre-task) and Rest 2 (post-task): Pearson’s r = 0.838, p≈0; Rest 1 and early learning: r = 0.765, p ≈ 0; Rest 2 and early learning: 0.761, p ≈ 0], where Rest 1 and Rest 2 were more similar (more highly correlated) than Rest 1 and early learning [Steiger’s test (a.k.a. Williams’ test and Meng’s test) for dependent correlations: t(30) = 20.508, p ≈ 0] and Rest 2 and early learning [t(30) = 21.777, p ≈ 0]. Notably, the correlation between Rest 1 and early learning did not differ significantly from that between Rest 2 and early learning [t(30) = 1.262, p = 0.207]. Thus, our prediction was confirmed for module allegiance matrices taken over all participants (Figure 7A. To bolster these findings, we calculated the same correlations for each pair of scans for each participant, testing the means (across participants) for the above relationships. This approach corroborated our findings, as the resting scans were again significantly more similar to each other than either of them was to early learning (Figure 7B). Among the behavioural subgroups, these findings were the case for FF, but not for SF and SS. Thus, the whole-group effect was driven by FF (Figure 7C).

**Fig. 7.**
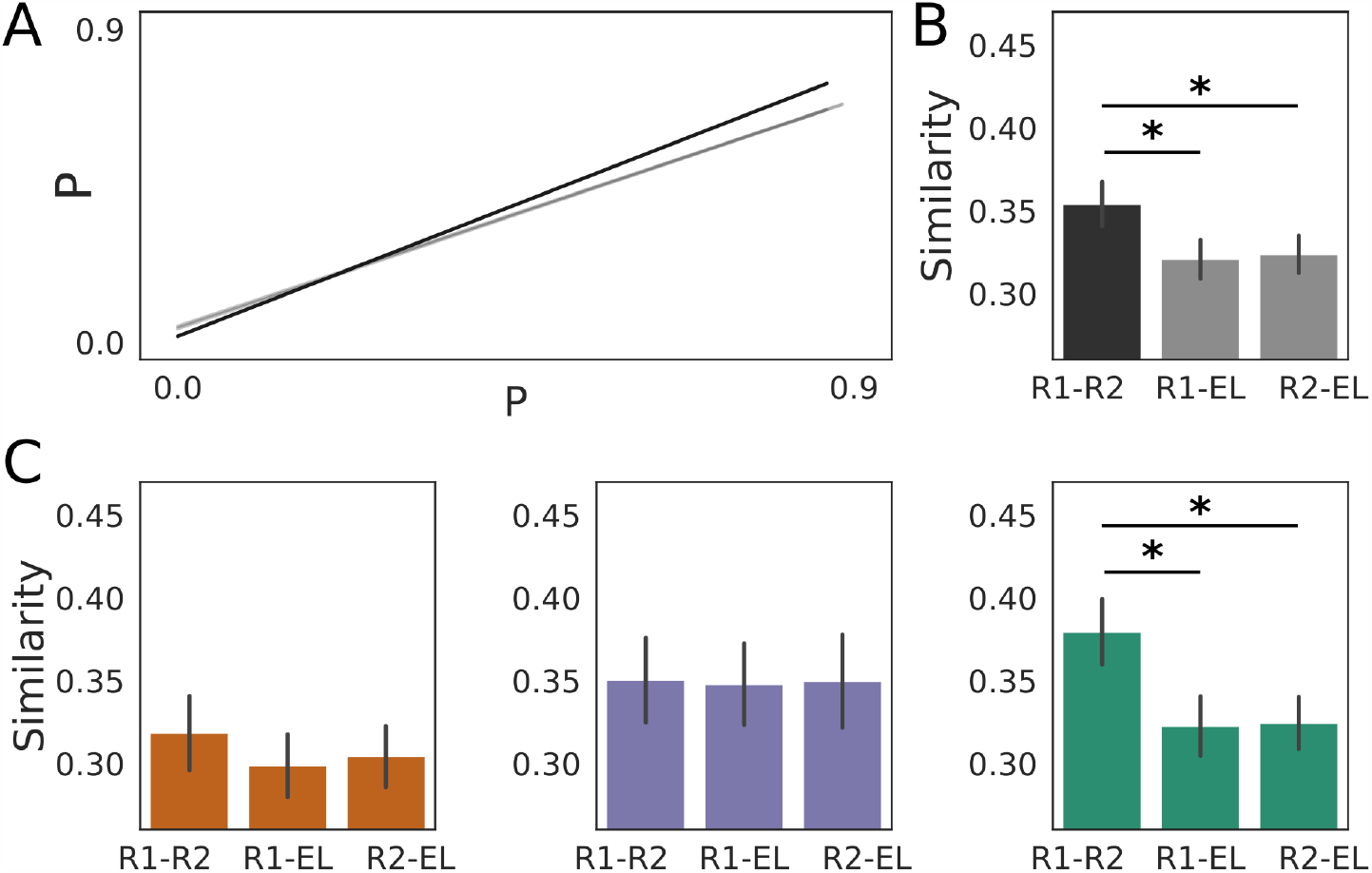
The organisation of dynamic, resting-state functional networks was more similar during pre-task (Rest 1) and post-task (Rest 2) rest than when either resting scan was compared to learning. **(A)** Linear fit to whole-group module allegiance probabilities (P, the probability that each of the 10,011 pairs of brain regions were in the same module; see text) for Rest 1 and Rest 2 (black, r = 0.838), Rest 1 and early learning (grey, r = 0.765) and Rest 2 and early learning (grey, r = 0.761). The grey lines are nearly indiscernible, as is the standard error in all fits. Fitted data points are omitted for clarity **(B)** Mean similarity (Pearson’s r) of module allegiance matrices across all participants for each pair of scans (R1: Rest 1; R2: Rest 2: EL: early learning). Rest 1 and Rest 2 were significantly more similar than Rest 1 and early learning [paired 1-tailed t-test: t(31) = 2.174, p = 0.019] and Rest 2 and early learning [R2-EL; t(31) = 1.918, p = 0.032], whereas the correlation between Rest 1 and early learning was not statistically different from Rest 2 and early learning [paired 2-tailed t-test: t(31) = -0.269, p = 0.790]. Note that our use of 1-tailed t-tests reflects our testing of predictions in panel ‘A’. **(C)** Mean similarity of each pair of scans for each behavioural subgroup (left, SS; middle, SF; right, FF). Among FF participants, module allegiance matrices from the resting scans were more similar than Rest 1 and early learning [paired 1-tailed t-test: t(14) = -2.112, p = 0.027] or Rest 2 and early learning [t(14) = -2.574, p = 0.011], whereas the resting scans’ correlations with learning were not significantly different [paired 2-tailed t-test: t(14) = 0.108, p = 0.542]. Among SF and SS participants, Rest 1 and Rest 2 were not significantly more similar than Rest 1 and early learning [paired 1-tailed t-test, SF: t(6) = -0.069, p = 0.474; SS: t(9) = -1.648, p = 0.067] or Rest 2 and early learning [SF: t(6) = -0.015, p = 0.494; SS: t(9) = -0.489, p = 0.318] and the correlation between Rest 1 and early learning was not statistically different from Rest 2 and early learning [paired 2-tailed t-test, SF: t(6) = 0.102, p = 0.539; SS: t(9) = 0.241, p = 0.592]. Single stars in panels B and C indicate statistical significance (p < 0.05).

## Discussion

Predictive biomarkers of cognition and learning are the subject of growing research interest (4, 8, 12, 13, 16, 18, 19). Within this body of work, the modularity of static functional networks at rest has been shown to be a promising measure for predicting neuropathologies and their responsiveness to treatment programs (16). Based on these findings, as well as on recent studies of the modular dynamics of cognition and learning (26), we asked whether dynamic modularity at rest would predict group differences on a task with well characterised behavioural variability, on which fast learning is widely believed to reveal the use of a cognitive strategy (27). First, we tested the hypothesis that modularity itself (the degree to which brain regions cluster together in functional subnetworks over time) would predict the learning profiles of participants who were fast learners on both days (FF), slow learners on both days (SS) and slow learners on the first day and fast learners on the second day (SF). At rest, prior to task performance, the modularity score *Q* was indeed associated with a fast learning profile on the upcoming task, distinguishing FF and SS participants (Figure 3). Second, we tested the hypothesis that the degree of coordination of modular reconfiguration would predict participants’ learning profiles, finding that FF and SS participants could be distinguished by the cohesiveness and disjointedness of modular changes at rest (Figure 4). Third, we tested the hypothesis that the recruitment and integration of functional networks derived during early learning, when the use of cognitive strategies is commonly believed to be most pronounced (30, 40), would predict participants’ learning profiles. We not only found that recruitment of the relearning network at rest was associated with fast adaptation, but we also found that its integration with (the regions of) the learning network was associated with slow adaptation, even though the learning network itself was not constructed until participants performed the task. Both measures distinguished the FF and SS subgroups (Figure 5). Overall, these findings are consistent with the hypothesis that modularity is not only a biomarker of clinical relevance, but is a more general biomarker of the propensity for cognition.

Of course, the propensity for cognition before an experimental task does not imply cognitive ability in a general sense, and we do not claim that measurements of dynamic modularity can (or should) be used to assess intelligence. For example, participants with higher modularity scores (or modularity-related statistics) may have been more alert and task-ready than their peers, owing to any number of factors, such the relative demands of their schedules, sleep, stress or caffeine consumption. Nonetheless, our predictive measures did distinguish fast learners from slow learners on the upcoming task, where fast learning is widely believed to be supported by the use of a cognitive strategy (27). Thus, we are comfortable claiming to have provided evidence that these measures reveal the conduciveness of participants’ wholebrain dynamics to the implementation of strategies. As such, our results speak to participants’ cognitive states, but they may not speak to cognitive traits (see Section below).

A related issue is the directionality of our resting-state predictions. We were able to distinguish pre-identified fast learners from slow learners by measures of modular dynamics at rest, but we did not assign participants to the subgroups based on these measures. Furthermore, all measures used here predicted fast learning relative to slow learning or *vice versa*, but it would be premature to propose a threshold for the use of these statistics, above or below which individuals may be classified as belonging to one group or another. Thus, despite the promise of our findings, it is unclear if measures of modular dynamics are suitable biomarkers for clinical diagnosis. Modular analysis of pooled resting-state data would help to determine the ‘normal range’ of modular measures.

### Our predictions were confirmed, but some results were surprising

Our hypotheses on *Q*, cohesion strength and disjointedness were borne out in the data, as each of these resting-state measures predicted learning outcomes in the expected manner. Specifically, greater *Q* (Figure 3) and cohesion strength (Figure 4A) were associated with fast learning, and greater disjointedness was associated with slow learning (Figure 4B). Our hypothesis that participants’ learning profiles would be predicted by resting-state recruitment and integration of task-based networks was also borne out, but we do not claim that our results on recruitment and integration were as expected. Other things being equal, we might have expected pre-task recruitment of the learning network to predict task performance, since this network was linked to fast learning on the same day (26). However, as described above, we were unable to derive the learning network during pre-task rest, indicating that its constituent regions began working together as a network during the task itself. Nevertheless, the more that its dispersed regions were integrated with the relearning network, the more likely participants were to be slow learners (Figure 5B). While this finding may not be considered surprising *per se*, we did not anticipate it. We were more surprised that recruitment of the relearning network predicted fast learning (Figure 5A) despite its inability to distinguish the subgroups during task performance until the next day (26). Since the networks were derived during the task on Day 1, it seems likely that all participants were using this network in some way, but their use of it did not differentiate the subgroups until Day 2, when (putatively) it was involved in recalling earlier strategies.

We were also surprised that the SS subgroup recruited the relearning network more strongly during post-task rest than pre-task rest (Figure 6). As such, SS participants’ organisation of their intrinsic networks became more like that of FF and SF participants, but this similarity with their faster-learning peers did not result in faster learning on the second day. Other things being equal, we might have expected the SF subgroup to show this sort of learning-induced change, since SF performance improved so dramatically on Day 2. In this regard, it is worth noting that we only investigated learning-induced changes in the measures for which we had pre-task hypotheses in the first place (see the Introduction), all of which predicted participants’ learning profiles. It is possible (perhaps likely) that other intrinsic network measures change after task performance, such that these changes would be informative about SF participants’ Day 2 improvement. Future work should address this possibility.

With respect to recruitment and integration more generally, it is worth reiterating that we derived the summary networks during the task (26) before quantifying their recruitment and integration during pre- and post-task rest. This approach contrasts with the more common one of measuring some aspect of pre-identified resting-state networks during task performance [*e*.*g*. Vatansever et al. (43), Gale et al. (44)]. Our approach is both philosophical and practical. The typical use of resting-state networks for task-based analyses leverages a common currency of sorts, assuming that the composition and properties of functional networks among a small number of people (on a variety of tasks) can be understood in terms of the composition and properties of intrinsic networks derived from a large number of people (45–48). This approach looks forward from a general predisposition to a specific outcome. The use of task-based networks for resting-state analyses is more tightly focused, looking backward from a specific outcome to a specific predisposition (among the same participants whose RSFC is under study). We believe these two approaches should be seen as complementary. Practically, our focus was the predictive value of modular dynamics during pre-task rest, so from this perspective, it made sense to use networks derived from dynamic modules linked to performance during the upcoming task.

### Networks derived at rest differed from those derived during learning

Our derivation of four summary networks from each of the first and second resting scans (Supplementary Materials Figure 2) was conspicuously different from the five networks derived during early learning (26). This difference was especially pronounced in relation to the learning network, which was no more than 20% similar to any network derived from either resting scan (quantified by the Jaccard index). This finding is consistent with the hypothesis that whole-brain cognitive control involves the ‘putting together’ of functional networks to suit the task at hand (26). Here, participants recruited (put together) the learning network during the SA task, pulling its regions from all of the four pre-task resting-state networks, except for the ‘restingstate relearning network’. Since all of these networks (rest and task) were derived from module allegiance matrices summarising modular dynamics in a given scan (41), we used these matrices to quantify the similarity of modular organisation in each pair of scans. In this regard, we found that pre- and post-task rest were more similar to one another than to early learning, which was (statistically) equally similar to both of these resting scans (Figure 7A-B). This whole-group finding was driven by the FF subgroup, as it was not the case for the SF and SS subgroups (Figure 7C). This result too is consistent with the hypothesis that cognitive control corresponds to recruitment of functional networks to suit the task at hand, since the FF subgroup was the only one whose behaviour implied a strategic approach to the task on Day 1. More generally, these findings suggest that the organisation of resting-state networks before and after tasks may be informative about the whole-brain mechanisms underlying task performance, including individual and group differences. Further work should investigate this intriguing possibility.

### Methodological considerations

At least two methodological considerations warrant further comment. Our first consideration is the very use of modularity and its statistics as predictive biomarkers of cognition. To begin with, modularity is a network measure. While early work on biomarkers demonstrated the predictive properties of individual brain regions, these findings were not robust across studies (16), suggesting that a common frame of reference across large groups of participants should be the first criterion for predictive biomarkers. If so, then the second criterion should be individual and group variation within that framework. RSFC provides this balance of competing factors (3). It is broadly accepted that cognitive phenomena are supported by largescale network interactions (49–51), so it makes sense to consider large-scale networks for cognitive biomarkers. Modularity confers many benefits to large-scale networks by balancing functional specialisation with integration over specialised subsystems (52). Most notably for the present work, modularity supports adaptability, enabling more efficient (53) and robust (54) learning. As such, modularity (quantified by *Q* here) appears to be an ideal biomarker for predicting the success of learning regimens for overcoming cognitive deficits, such as those associated with brain injury (20) and old age (21). Nonetheless, the relationship between *Q*, modular reconfiguration and subnetwork recruitment is not straightforward. It is plausible that modularity is a wholebrain trait that requires coordination among regions to produce qualitative changes in whole-brain states, supporting the recruitment of functional networks. This characterisation of modular dynamics is consistent with the association of deeper (shallower) states of unconsciousness with more disjointed (cohesive) modular reconfiguration (55), but further research is needed to understand such large-scale network dynamics. Whole-brain models and formal analysis of their emergent dynamics are an exciting direction in this regard (56).

A second consideration is our use of PC1 to identify predictive relationships between neural variables and participants’ learning profiles. While our use of PC1 in this way is well justified (26), this correlation-based approach will only identify neural variables of interest where SF is sandwiched between FF and SS (or is roughly equal to either of them) since PC1 corresponds to mean early error across days (Figure 2D). For *Q*, cohesion strength and disjointedness, this issue did not matter, as our hypotheses were confirmed. For recruitment and integration, however, we looked for relationships between PC1 and the recruitment of each of five networks, and between PC1 and the integration of the relearning network with the other networks (both sets of tests were corrected for multiple comparisons, described in the Results). Had we looked for predictive relationships between neural variables and PC2 (which corresponds to savings, Figure 2D) we may have identified differences between SF and the other two subgroups, given SF’s pronounced improvement in learning performance on Day 2. Confirmation of each of our hypotheses clearly differentiated the FF and SS subgroups, which were comprised of unambiguously fast (explicit) and slow (implicit) learners. The SF subgroup is (and remains) something of a curiosity. We leave this consideration to a follow-up study.

### Summary and conclusions

Modularity is a promising predictive biomarker of cognitive pathologies and their responsiveness to clinical programs (16). Where most earlier work focused on static modularity (20–22), quantifying the subnetwork composition of functional networks that average out the dynamics of resting-state activity, we featured these dynamics by quantifying the subnetwork composition of functional networks over time [see also (57)]. By characterising the time-evolution of this composition, we have demonstrated that standard measures of dynamic modularity (26, 39, 41, 55) at rest are able to predict learning profiles on a task with well-characterised individual differences (24–26). These differences are widely believed to reveal the degree to which participants take a strategic (cognitive) approach to the task (27), so our results highlight modularity as a more general cognitive biomarker. They further highlight the promise of dynamic modularity in the clinical domain, with a view to diagnosis and treatment of cognitive pathologies.

## Methods

### Experimental design and statistical analysis

40 righthanded individuals between the ages of 18 and 35 (mean = 22.5, standard deviation = 4.51; 13 males) participated in the study and received financial compensation for their time. Data from 8 participants were excluded, due to either head motion in the MRI scanner (N=4; motion greater than 2 mm or 2^*°*^ rotation within a single scan) or missing volumes in a large portion of one of the task scans (N=4), leaving 32 participants in the final analysis. Right handedness was assessed with the Edinburgh handedness questionnaire (58). Participants’ written, informed consent was obtained before commencement of the experimental protocol. The Queen’s University Research Ethics Board approved the study, which was conducted in accordance with the principles outlined in the Canadian Tri-Council Policy Statement on Ethical Conduct for Research Involving Humans and the principles of the Declaration of Helsinki (1964).

Experimenters were not blind to testing. Statistical significance was defined by *α* = 0.05. In all figures, error bars indicate *±* 1 standard error of the mean (SEM). All statistical tests were two-tailed. FDR correction for family-wise error rate used the method by (59).

### Apparatus

In the scanner, participants performed hand movements that were directed towards a target by applying a directional force onto an MRI-compatible force sensor (ATI technologies) using their right index finger and thumb. The target and cursor stimuli were rear-projected with an LCD projector (NEC LT265 DLP projector, 1024 x 768 resolution, 60 Hz refresh rate) onto a screen mounted behind the participant. The stimuli on the screen were viewed through a mirror fixed to the MRI coil directly above participants’ eyes, thus preventing participants from being able to see the hand. The force sensor and associated cursor movement were sampled at 500 Hz.

### Procedure

The experiment used a well-established VMR task (36) to probe sensorimotor adaptation (Figure 1A). At the start of each trial, the cursor (20-pixel radius) appeared in a central position (25-pixel radius). A white target circle (30-pixel radius) was presented on a black screen, appearing at one of eight locations (0, 45, 90, 135, 180, 225, 270, 315^*°*^) on an invisible ring around the central position (300-pixel radius) and filled in (white) following a 200 ms delay.

Once filled in, participants applied a brief directional force to the sensor (threshold of 1.5 N), which launched the cursor toward the target. Target location was drawn at random from the above locations without replacement, in bins of 8 trials. Once launched, the cursor travelled the 300-pixel distance to the ring over a 750 ms period (with a bell-shaped velocity profile) before becoming stationary at the ring to provide participants with end-point error feedback. If the cursor overlapped the target to any extent, the target would become green, signifying a ‘hit’. Each trial was separated by 4 s and within this period, participants had 2.6 s from target presentation to complete the trial (including the 200 ms target delay, participants’ reaction time and the 750 ms cursor movement; any remaining time was allotted to the end-point error feedback). At 2.6 s, the trial was ended, the screen blanked, and the data saved, and participants briefly waited for the next trial to begin. Reaction times were not emphasised. On trials in which the reaction time exceeded 2.6 s, the trial ended, and the participant waited for the next trial to begin. These discarded trials were rare (0.56% across all trials and participants) and were excluded from behavioural analyses. They were kept in the neuroimaging analysis due to the continuous nature of the fMRI task and our focus on functional connectivity.

During each testing session, 120 baseline trials (15 bins of 8 trials) were completed without a rotation of the cursor. Following these trials, 320 learning trials (40 bins of 8 trials) were completed, during which a 45^*°*^ clockwise rotation of the cursor was applied. The baseline and learning trials were completed during one continuous fMRI scan. Following this scan, conditions were restored to baseline (*i*.*e*. no rotation of the cursor) in a separate scan and participants performed 120 washout (unlearning) trials. These washout trials allowed us to probe participants’ rate of relearning 24 hours later (and thereby their savings). In addition to these VMR-related task components, we interspersed three 6-minute resting fMRI scans before, between and after VMR learning and washout. During resting scans, participants were instructed to rest with their eyes open, while fixating a central cross presented on the screen. The total testing time was 75 minutes on each testing day.

### MRI acquisition

Participants were scanned using a 3-Tesla Siemens TIM MAGNETOM Trio MRI scanner located at the Centre for Neuroscience Studies, Queen’s University (Kingston, Ontario, Canada). Functional MRI volumes were acquired using a 32-channel head coil and a T2*-weighted single-shot gradient-echo echo-planar imaging (EPI) acquisition sequence (time to repetition (TR) = 2000 ms, slice thickness = 4 mm, in-plane resolution = 3 mm x 3 mm, time to echo (TE) = 30 ms, field of view = 240 mm x 240 mm, matrix size = 80 x 80, flip angle = 90°, and acceleration factor (integrated parallel acquisition technologies, iPAT) = 2 with generalised auto-calibrating partially parallel acquisitions (GRAPPA) reconstruction. Each volume comprised 34 contiguous (no gap) oblique slices acquired at a 30° caudal tilt with respect to the plane of the anterior and posterior commissure (AC-PC), providing whole-brain coverage of the cerebrum and cerebellum. A T1-weighted ADNI MPRAGE anatomical was also collected (TR = 1760 ms, TE = 2.98 ms, field of view = 192 mm x 240 mm x 256 mm, matrix size = 192 x 240 x 256, flip angle = 9°, 1 mm isotropic voxels). For each resting scan, 180 imaging volumes were collected. For the baseline and learning epochs, a single, continuous scan was collected of 896 imaging volumes duration. For the washout scan, one scan of 256 imaging volumes was collected. Each scan included an additional 8 imaging volumes at both the beginning and end of the scan.

On Day 1, a separate practice session was carried out before the actual fMRI experiment to familiarise participants with the apparatus and task. This session involved performing 60 practice baseline trials. The fMRI testing session for each participant lasted 2 hours and included set-up time (20 minutes), practice (10 minutes), one high-resolution anatomical scan (8 minutes), two DTI scans (one in the AP direction and the other in the PA direction; 10 minutes), a resting scan (6 minutes), a baseline and rotation scan (30 minutes), a resting scan (6 minutes), a washout scan (9 minutes), and a final resting scan (6 minutes). In the present paper, we only consider fMRI data from the first and second resting scans, and the baseline-learning scan, on the first day.

### MRI preprocessing

Preprocessing was performed with fMRIPrep 1.4.0 [(60, 61) RRID:SCR_016216].

#### Anatomical data preprocessing

T1w scans in each session were corrected for intensity non-uniformity (INU) with N4BiasFieldCorrection (62), distributed with ANTs 2.2.0 (63). The T1w-reference was then skull-stripped with a Nipype implementation of the antsBrainExtraction.sh workflow (from ANTs), using OASIS30ANTs as target template. Brain tissue segmentation of cerebrospinal fluid (CSF), white-matter and grey matter was performed on the brain-extracted T1w using fast [FSL 5.0.9 (64)]. A T1w-reference map was computed after registration of the individual T1w scans (after INU-correction) using mri_robust_template [FreeSurfer 6.0.1 (65)]. Brain surfaces were reconstructed using recon-all [FreeSurfer 6.0.1 (66)], and the estimated brain mask was refined with a custom variation of the method to reconcile ANTs-derived and FreeSurfer-derived segmentations of the cortical grey matter of Mindboggle (67). Volume-based spatial normalisation to FSL’s MNI ICBM 152 non-linear 6th Generation Asymmetric Average Brain Stereotaxic Registration Model [(68); TemplateFlow ID: MNI152NLin6Asym] was performed by nonlinear registration with antsRegistration (ANTs 2.2.0), using brain-extracted versions of both T1w reference and the T1w template.

#### Functional data preprocessing

For each BOLD run (for each participant), a reference volume and its skull-stripped version were generated using a custom methodology with fMRIPrep. The BOLD reference was then co-registered to the T1w reference using bbregister (FreeSurfer), implementing boundary-based registration (69). Co-registration was configured with nine degrees of freedom to account for distortions remaining in the BOLD reference. Head-motion parameters with respect to the BOLD reference (transformation matrices, and six corresponding rotation and translation parameters) were estimated before any spatiotemporal filtering using mcflirt [FSL 5.0.9 (70). BOLD runs were slicetime corrected using 3dTshift from AFNI 20160207 [(71) RRID:SCR_005927]. The BOLD time-series were resampled into standard space (MNI152NLin6Asym), generating spatially-normalised, preprocessed BOLD runs. A reference volume and its skull-stripped version were generated using a custom methodology with fMRIPrep. Several confounding time-series were calculated based on the preprocessed BOLD. The six head-motion estimates calculated in the correction step were included as confounding time series, along with the temporal derivatives and quadratic terms. A set of physiological regressors were extracted to allow for component-based noise correction (72). Principal components were estimated after high-pass filtering the preprocessed BOLD time-series (using a discrete cosine filter with 128s cut-off) based on anatomical masks (aCompCor). We used aCOMPCOR because it has proven to be one of the most consistent pipelines for mitigating the problem of motion across performance indices with dynamic functional connectivity analyses (73). Components were calculated within the intersection of the aforementioned mask and the union of CSF and white matter masks calculated in T1w space, after their projection to the native space of each functional run (using the inverse BOLD-to-T1w transformation). Components were also calculated separately within the white matter and CSF masks. *k* components with the largest singular values were retained, and the remaining components were dropped from consideration. All resamplings were performed with a single interpolation step by composing all the relevant transformations (*i*.*e*. head-motion transform matrices, susceptibility distortion correction when available, and co-registrations to anatomical and output spaces). Gridded (volumetric) resamplings were performed using antsApplyTransforms (ANTs), configured with Lanczos interpolation to minimise the smoothing effects of other kernels (74). Non-gridded (surface) resamplings were performed using mri_vol2surf (FreeSurfer).

Many internal operations of fMRIPrep use Nilearn 0.5.2 [(75) RRID:SCR_001362], mostly within the functional processing workflow. For more details of the pipeline, see the section corresponding to workflows in fMRIPrep’s documentation.

#### Region of interest signal extraction

Region of interest (ROI) extraction and nuisance regression were performed using Nilearn 0.5.2 (75). For each participant, the raw BOLD time series for each voxel inside a given ROI was isolated using the Schaeffer-100 atlas for cortical areas (76), the Harvard-Oxford atlas for subcortical areas (77–80), and the SUIT cerebellar atlas for the cerebellum (81). The mean time course of each ROI was high-pass filtered (cutoff = 0.01Hz), temporally detrended and z-scored across each run, in order to ensure that all voxels had the same scale of activation after preprocessing. In order to account for motion confounds and physiological noise, the above-described nuisance regressors were included, *i*.*e*. participants’ six motion parameters and their derivatives, squares, and the squares of the derivatives (24 total) and the top six aCompCor regressors. Note that nuisance regression was performed orthogonally to temporal filters, so as to avoid reintroducing high-frequency noise from the regressors (82).

### Data Analysis

#### Behavioural analyses

To assess performance on the task, we calculated the angle of the cursor relative to the target at the moment the cursor reached the target distance (or the nearest data point if this value could not be used). This calculation was performed by subtracting the target angle from an angle derived from the horizontal and vertical position at a 300-pixel distance from the starting point. The median endpoint cursor error for the bin of 8 trials (1 trial for each target) was extracted (binned median error) and the mean binned median error for early trials (bins 1-3; 24 trials total) and late trials (bins 38-40; 24 trials total) were calculated to yield ‘early’ and ‘late’ error measures for all participants on each day. Savings was measured by subtracting the early error on Day 2 from the early error on Day 1 (32). To determine whether distinct subgroups of learners were present in our sample, k-means clustering (83) was performed across participants using these five learning measures (early error on Day 1, early error on Day 2, late error on Day 1, late error on Day 2, and savings). We ran k-means until convergence 1000 times with random initialization for all integers in 2 ≤ k ≤ 9 (maximum of 10,000 iterations of the algorithm for each initialization). The solution with the lowest sum of Euclidean distances between the data points and their assigned centroids was chosen for each value of k, and each of these solutions was evaluated by the Silhouette (84) and Calinski-Harabasz (85) cluster evaluation indices. The solution corresponding to k = 3 had the highest score according to both indices [see (26)]. Results were identical under Euclidean and Manhattan distances measures. We tested this clustering solution for statistical significance by randomly drawing from a multivariate normal distribution with the same means and covariance as the data (1000 iterations, one draw for each participant), running k-means for each iteration in the same way as above. We recorded the highest value of the Silhouette and Calinski-Harabasz indices (over all k) for each iteration, where the probability of falsely rejecting the null hypothesis (for each evaluation index) was equated with the proportion of iterations with a higher score than our clustering solution.

#### Temporal modularity of functional networks

Following earlier work (38), the maximum overlap discrete wavelet transform was used to decompose the time series for each ROI into wavelet coefficients in the range of 0.12 - 0.25Hz (Haar wavelet family, scale 1). The decomposed time series were divided into windows of w=64 seconds (32 imaging volumes; corresponding to 2 bins of 8 trials; see above) where contiguous windows overlapped by 1 bin, *i*.*e*. by 50% (55, 86). We constructed functional networks in each time window by calculating the magnitude squared spectral coherence between each pair of brain regions and taking the mean coherence over the specified range of frequencies (38). All coherence values less than the 95th percentile of a null model were set to zero. For each of baseline, learning and unlearning, the null model was constructed over 10,000 iterations by selecting two regions and start times at random, shuffling their contents over window w, and calculating their coherence in the same way as for the original time series. This approach produced multislice networks with a mean sparsity of ∼ 85%. We determined the modular composition of the resulting multislice networks with a generalised Louvain method for time-resolved clustering (87). This algorithm was repeated 100 times, resulting in 100 clustering solutions (a.k.a. partitions), each of which maximised a modularity ‘quality’ function (37). On each repetition, we initialised the algorithm at random (uniform distribution) and iterated until a stable partition was found, *i*.*e*. the output on iteration *n* served as the input on iteration *n* + 1 until the output matched the input (37, 38), where nodes were assigned to modules at random from all assignments that increased the quality function, and the probability of assignment was proportional to this increase. We used the standard spatial and temporal resolution parameters *γ* = 1 and *ω* = 1 respectively, and the standard null model by Newman and Girvan (88) for the quality function (23, 37).

#### Module allegiance matrix

We constructed a matrix *T*, where the elements *T*_*i,j*_ refer to the number of times regions *i* and *j* were assigned to the same module over all time slices, partitions and participants during early learning [wavelet scale 2 (26)] and (separately) the first resting scan. We then constructed the module allegiance matrix *P* = (1*/C*)*T*, where *C* is the total number of time slices in all of these partitions. Values that did not exceed those of a null model were set to 0 (89).

#### Clustering of the module allegiance matrix with non-negative matrix factorisation

To derive summary networks from the module allegiance matrix, we used non-negative matrix factorisation [SymNMF; (90, 91)], which decomposes a symmetric matrix *Y* as *Y* = *AAT*, where *A* is a rectangular matrix containing a set of nonnegative factors that minimise a sum-of-squares error criterion. SymNMF has previously shown good performance in community detection problems (90, 92) and has the benefit of producing a natural clustering solution by assigning each observation to the factor on which it loads most strongly. We used the Newton-like Sym-NMF algorithm described by Kuang et al. (90), where for each number of factors between 2 and 15, we fit the model using 250 random initializations of the factor matrix H, with entries sampled from a uniform distribution on the interval [0, 1]. We then computed, for each rank, the average root mean squared reconstruction error, the dispersion coefficient (93) and the cophenetic correlation (94). The latter two measures quantify the consistency of the clustering solutions returned by different initialisations. We judged that a rank 5 solution was acceptable, as it explained over 80% of the variance in the observed data, it was acceptable by both the dispersion and cophenetic correlation criteria, and the generation of additional clusters provided little improvement in explained variance. We then selected the rank 5 solution with the lowest reconstruction error, and generated a clustering solution by assigning each brain region to the factor with the highest loading.

## ACKNOWLEDGEMENTS

This work was supported by the Canadian Institutes of Health Research and the Natural Sciences and Engineering Research Council of Canada.

