## Supplementary Materials for "Whole-brain modular dynamics at rest predict sensorimotor learning performance"

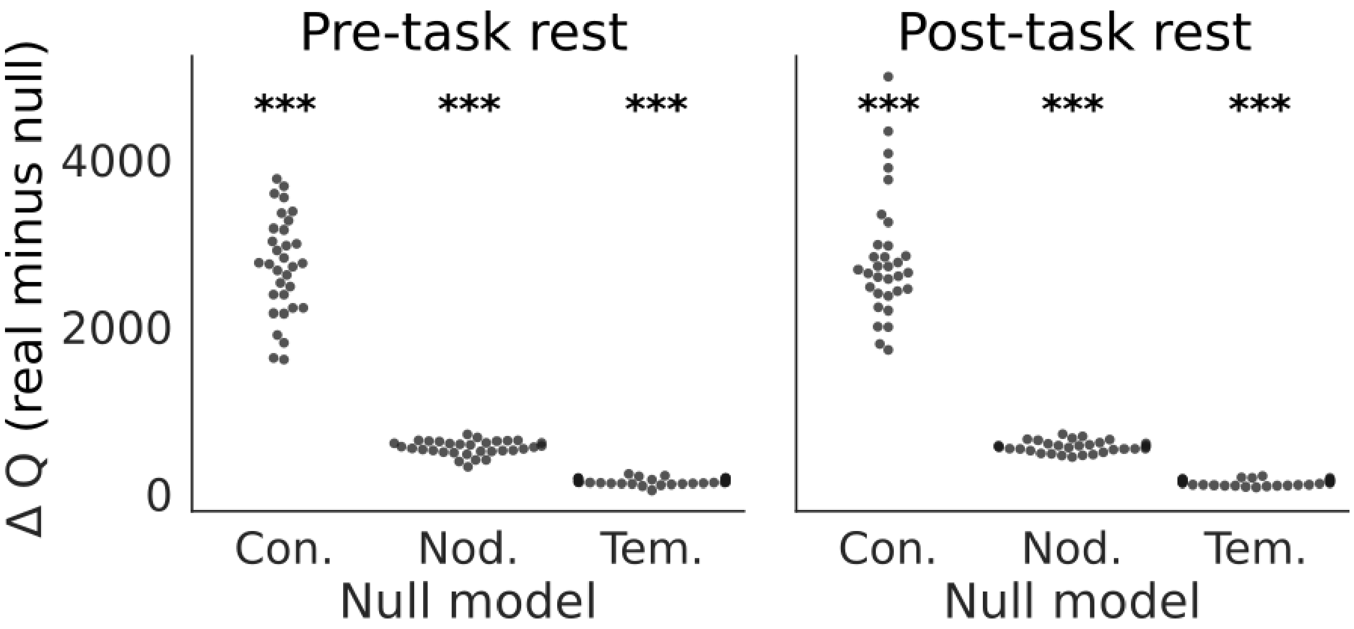


**Fig. 1. Temporal networks were significantly modular at rest, before and after learning.** Scatter plots show the difference ($\Delta Q$) in the modular quality function $Q$ between real and permuted networks (real minus permuted) for connectional (Con.), nodal (Nod.) and temporal (Tem.) null models (1) during pre-task (left) [connectional: $t(31) = 26.329$, $p = 8.911e-23$; nodal: $t(31) = 35.372$, $p = 1.261e-26$; temporal: $t(31) = 21.158$, $p = 5.525e-20$] and post-task (right) [connectional: $t(31) = 21.805$, $p = 2.298e-20$; nodal: $t(31) = 45.960$, $p = 4.365e-30$; temporal $t(31) = 21.272$, $p = 4.721e-20$] rest. Each dot corresponds to a participant. Jitter is for visual clarity. As in the main text, three stars indicate $p < 1e-3$.

### Supplementary Note 1: Resting-state functional networks were significantly modular

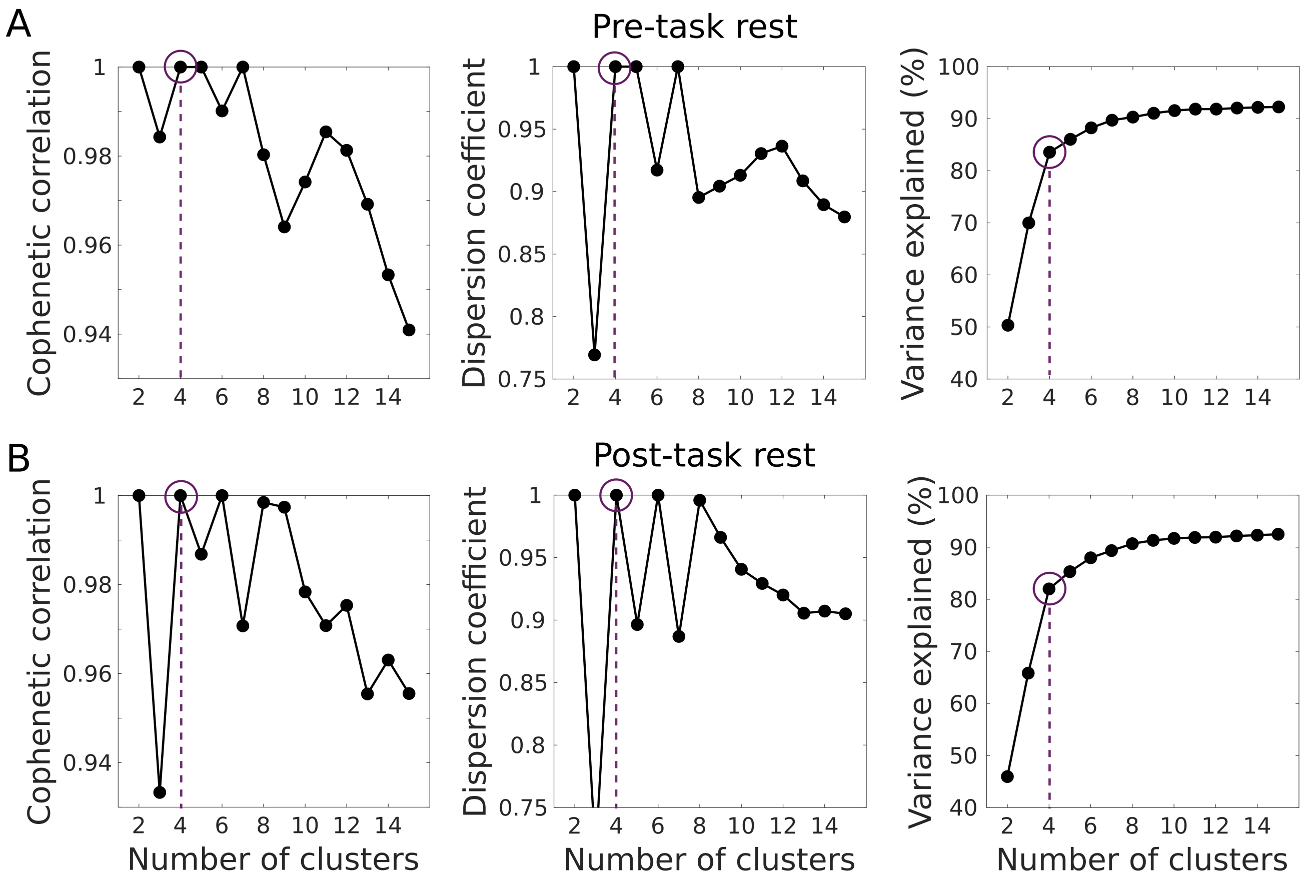


**Fig. 2. Clustering diagnostics for symmetric non-negative matrix factorization of pre- and post-task module allegiance matrices.** Symmetric NMF was performed using 250 random initializations for factors 2 to 15. Regions were then assigned to a cluster corresponding to the factor with the greatest loading. For each number of factors, we computed the cophenetic correlation, dispersion, and explained variance. Dashed vertical line in each panel indicates the chosen solution of 4 clusters for pre-task (A) and post-task (B) rest.

### Supplementary Note 2: Clustering diagnostics identified four resting-state networks during pre- and post-task rest
